# Single Cell Cortical Bone Transcriptomics Defines Novel Osteolineage Gene Sets Altered in Chronic Kidney Disease

**DOI:** 10.1101/2022.07.30.502024

**Authors:** Rafiou Agoro, Intawat Nookaew, Megan L. Noonan, Yamil G. Marambio, Sheng Liu, Wennan Chang, Hongyu Gao, Lainey M. Hibbard, Corinne E. Metzger, Daniel Horan, William R. Thompson, Xiaoling Xuei, Yunlong Liu, Chi Zhang, Alexander G. Robling, Lynda F. Bonewald, Jun Wan, Kenneth E. White

**Affiliations:** Department of Medical and Molecular Genetics, Indianapolis, IN 46202, USA; Department of Biomedical Informatics, University of Arkansas for Medical Sciences, AR 72205, USA; Center for Medical Genomics, Indiana University School of Medicine, Indianapolis, IN 46202, USA; Department of Electrical and Computer Engineering, Purdue University, Indianapolis, IN 46202, USA; Department of Anatomy, Cell Biology and Physiology, Indianapolis, IN 46202, USA; Indiana Center for Musculoskeletal Health, Indiana University, Indianapolis, IN 46202, USA; Department of Medicine/Nephrology; Indiana University School of Medicine, Indianapolis, IN 46202

**Keywords:** Bone, scRNAseq, Osteoblast, Osteocyte, Chronic Kidney Disease

## Abstract

Due to a lack of spatial-temporal resolution at the single cell level, the etiologies of the bone dysfunction caused by diseases such as normal aging, osteoporosis, and the metabolic bone disease associated with chronic kidney disease (CKD) remain largely unknown. To this end, flow cytometry and scRNAseq were performed on long bone cells from Sost-cre/Ai9^+^ mice, and pure osteolineage transcriptomes were identified, including novel osteocyte-specific gene sets. Clustering analysis isolated osteoblast precursors that expressed *Tnc*, *Mmp13,* and *Spp1,* and a mature osteoblast population defined by *Smpd3*, *Col1a1*, and *Col11a1*. Osteocytes were demarcated by *Cd109*, *Ptprz1, Ramp1, Bambi, Adamts14*, *Spns2, Bmp2*, *WasI,* and *Phex*. We validated our *in vivo* scRNAseq using integrative *in vitro* promoter occupancy via ATACseq coupled with transcriptomic analyses of a conditional, temporally differentiated MSC cell line. Further, trajectory analyses predicted osteoblast-to-osteocyte transitions via defined pathways associated with a distinct metabolic shift as determined by single-cell flux estimation analysis (scFEA). Using the adenine mouse model of CKD, at a time point prior to major skeletal alterations, we found that gene expression within all stages of the osteolineage was disturbed. In sum, distinct populations of osteoblasts/osteocytes were defined at the single cell level. Using this roadmap of gene assembly, we demonstrated unrealized molecular defects across multiple bone cell populations in a mouse model of CKD, and our collective results suggest a potentially earlier and more broad bone pathology in this disease than previously recognized.

## Introduction

Osteocytes are derived from osteoblasts through a dynamic, spatio-temporal process regulated within cortical bone [1]. The exposure of these cells to the bloodstream through dendrites allows osteocytes to play key endocrine functions, including sending and receiving signals to vascularized organs such as the kidney, among others. A complete identification of the osteocyte and osteocyte precursor transcriptomes is a critical need towards understanding the molecular pathways of osteolineage differentiation, as well as to specify the contributions of osteoblast/osteocyte genes to musculoskeletal disease.

Common bone pathologies are associated with osteocyte dysfunction, including aging [2], osteoporosis [3], and chronic kidney disease (CKD) [4]. CKD is a worldwide public health problem with an estimated 5-10 million lives lost each year [5]. Endocrine changes due to loss of renal function lead to increased bone production of fibroblast growth factor-23 (FGF23), which acts on the kidney to reduce circulating 1,25(OH)_2_ vitamin D (1,25D) concentrations. These actions in turn cause hypocalcemia-mediated compensatory elevations in PTH and subsequent bone loss due to increased osteoclastic activity [6]. Indeed, fracture is the most important clinical outcome of the CKD bone disorder, with an estimation that CKD patient hip and spine fracture rates range between 2- to 4-fold greater than the general population [7], markedly increasing patient morbidity and mortality [8].

Several studies have provided initial insight into novel bone cell bioactivity using the analysis of bulk sequencing of cortical bone [9] and single cell analyses of combined cortical bone and calvaria cells [10]. These approaches have led to an increased understanding of transitional states of the skeletal cell populations in diseases typically associated with progressive loss of bone mass, including osteoporosis and aging [11]. Additionally, the previous studies identified novel osteoblast and osteocyte genes, providing key insight into new roles of these loci for a potentially broader set of bone diseases. However, the full spectrum of osteoblast and osteocyte genes in bone cell populations specifically affected during CKD remain understudied, hampering the ability to target cell lineages towards improved bone health and patient outcomes. Thus, the nature of osteolineage cells affected during CKD, as well as the molecular mechanisms and spatial transcriptional reprogramming that contribute to bone loss remain to be determined.

Herein, we developed a successful workflow to enrich cortical bone osteoblast and osteocyte cell populations using flow cytometry followed by single cell RNAseq (scRNAseq). Our transcriptomic data detected two osteoblast populations, as well as novel osteocyte-specific genes, and also revealed cell-specific metabolic profiles. This approach identified previously unrecognized transcriptional reprogramming prior to major bone ultrastructural changes in a mouse model of CKD. Our collective results suggest that the endocrine bone disease arising from CKD affects multiple cell populations in parallel and thus emerging treatments for CKD may be more effective by targeting a broader set of cell lineages.

## Methods

### Sost-CreERT2-Cre+/Ai9 mice

Animal protocols and studies were approved by the Indiana University Institutional Animal Care Committee and conform to NIH guidelines [12]. Male and female Sost-CreERT2-Cre+ mice were crossed with the ‘Ai9’ mouse in which the recombination event causes activation of tdTomato fluorescent protein to generate Sost-CreERT2 Cre+/Ai9 [13]. Both males and females were included in the scRNAseq study. For the sequencing experiments, with the goal to isolate cells in their native state without confounding effects on bone, due to the presence of basal Cre activation in Sost-CreERT2-Cre+ mice [13], six Sost-CreERT2/Ai9 mice at eight weeks of age were directly sacrificed without tamoxifen injection, and long bones (femurs, tibiae, and humeri) were collected and pooled, followed by a sequential bone digestion (see below).

### Preparation of mouse long bones for osteoblast/osteocyte isolation

Isolated long bones were placed in 100 mm petri dishes containing αMEM with 10% penicillin and streptomycin. Any remaining muscle and connective tissue from the bones were removed and the periosteum was scraped away using a scalpel. The bones were then washed, epiphyses removed, and the marrow flushed using a 27G syringe. The flushed bones were then cut in half lengthwise and into 1 to 2 mm lengths using a scalpel and briefly washed with Hank’s Balanced Salt Solution (HBSS; Hyclone).

### Bone digestion and isolation of osteoblasts and osteocytes

The long bone pieces were digested as previously described [14]. In brief, bones were alternately immersed in warmed collagenase type IA (2 mg/mL; Sigma) and EDTA (5 mM), then incubated at 37°C for 25 min on a rotating shaker (∼200 rpm). A total of 9 digestions was performed. The extracted bone cells were then centrifuged at 600 rpm for 15 minutes at 4°C, washed 3 times in αMEM supplemented with 10% FBS and the pellet resuspended in αMEM supplemented with 10% FBS. TdTomato^+^ cells were then sorted using Fluorescence-activated cell sorting (FACS) Aria Fusion (BD Biosciences) with standard Phycoerythrin (PE) Texas Red gating. After cell isolation, cell viability was assessed using Trypan Blue staining; more than 90% of sorted cells were viable and these cells were immediately processed for scRNAseq.

### CKD mouse models

Male and female C57BL/6J mice were purchased from the Jackson Laboratory and housed at least one week in the Indiana University School of Medicine Laboratory Animal Resource Center (LARC) for acclimatization prior to the start of experiments, according to previous protocols [15]. The adenine diet model was used to induce CKD in male and female mice. At 8 weeks of age, mice were fed a control casein-based diet (0.9% phosphate and 0.6% calcium, TD.150303 Envigo) for 2 weeks, or CKD-causing diet with 0.2% added adenine (TD.160020; Envigo) for 2 or 4 weeks. Diets and water were provided ad libitum.

### Osteocyte and osteoblast cell counting in cortical and trabecular bone

Femurs from mice fed with the casein control or adenine-CKD diets for 2 or 4 weeks were fixed in 4% paraformaldehyde for 24 hours. The samples were then decalcified in a solution that contains 1.2% neutral buffered formalin and 10% EDTA for 12-days at 4°C on a rocker platform. The decalcified solution was changed three times over the decalcification period. The femurs were then washed in water and stored in 70% ethanol at 4°C. For histological analysis, the femurs were dehydrated overnight and embedded in paraffin. Sections of 5 µm of each sample were stained with hematoxylin eosin (H&E) for light microscopy. For counting osteocytes in cortical bone, a blinded experiment was conducted and an area of cortical bone of approximately ∼550 mm^2^ in the midshaft was analyzed for cell numbers followed with a normalization to bone surface. Analyses were performed using BIOQUANT imaging (BIOQUANT Image Analysis, Nashville, TN). Trabecular osteocytes were analyzed and normalized to trabecular bone area in the distal femur excluding endocortical surfaces and primary spongiosa. Osteoblast numbers were normalized to trabecular bone surface. Three mice per condition were used for these experiments.

### Micro-computed tomography (μCT)

Paraformaldehyde fixed femora from CKD and healthy mice were scanned, reconstructed, and analyzed as previously described [16]. Femurs were scanned at 10-μm resolution, 55-kV peak tube potential and 8W. Standard output parameters related to cortical bone mass, geometry, and architecture were measured as reported [17].

### Single cell library preparation

We applied a single cell master mix with lysis buffer and reverse transcription reagents according to the Chromium Single Cell 3’ Reagent Kits V3 User Guide, CG000183 Rev A (10X Genomics, Inc.). This was followed with cDNA synthesis and library preparation according to standard 10X Genomics methods; all libraries were sequenced on an Illumina NovaSeq6000 platform in paired-end mode (28bp + 91bp).

### Data processing/bioinformatic analyses

The 10x Genomics Cellranger (v. 6.0.0) pipeline was used to demultiplex raw base call files to FASTQ files and reads were aligned to the mm10 murine genome using STAR [18]. The Cellranger computational output was then analyzed in R (v 4.0.2) using the Seurat package v. 4.1.0 4 [19]. Seurat objects were created, and the top principal components were used to perform unsupervised clustering analysis and visualized using UMAP dimensionality reduction. Using the Seurat package, annotation and grouping of clusters by cell type was performed manually by inspection of differentially expressed genes using the MAST method [20] for each cluster, based on canonical marker genes in the literature. The selected markers were visualized on UMAP coordinates as gene expression density using the R package Nebulosa [21]. To perform the pseudotime analysis on the integrated Seurat object, cells were divided into individual gene expression data files organized by previously defined cell types. R package Monocle v3 [22] was used for dataset analysis and outputs were obtained detailing the pseudotime cell distributions for each cell type. Positional information for the Monocle plot was used to subset and color cells for downstream analyses [22]. Ingenuity Pathway Analysis (IPA) was used to predict the statistically significant canonical pathways regulated in osteolineage cells (pre-osteoblasts, osteoblasts and osteocytes).

### scFEA

The python package v1.2 of single cell flux estimation analysis (scFEA) was applied to estimate cell-wise metabolic flux rate against the whole human metabolic map using the generated mouse scRNAseq data [23]. Default parameters were utilized and statistical significance of the differences in metabolic flux between cell groups was assessed by Mann-Whitney test.

### Culture of the MPC cell line

A conditionally immortalized mesenchymal stem cell line, Murine progenitor cells clone 2 (MPC2) [24], was cultured in αMEM (Invitrogen, Thermo-Fisher Scientific) supplemented with 10% fetal bovine serum (FBS; Hyclone), 25 mM L-glutamine, and 25 mM penicillin-streptomycin (Sigma-Aldrich, St. Louis, MO, USA) at 33°C and 5% CO_2_ to proliferate. Cells were plated at a density of 1.0×10^5^ cells per well in 6-well plates and incubated overnight before being transferred to a 37°C incubator for osteogenic differentiation by culturing in maintenance media supplemented with 4 mM beta-glycerophosphate and 50 μg/mL ascorbic acid. Cells were differentiated for 0-4 weeks with this ‘osteogenic media’, which was changed every 2-3 days.

### Bulk mRNA sequencing

MPC2 cells were differentiated for 3 weeks in osteogenic media or plated in an undifferentiated state at 33°C (see cell culture methods above). Total RNA was extracted and evaluated for its quantity and quality using an Agilent Bioanalyzer 2100; 100 ng of total RNA was used for the cDNA libraries. Library preparation included mRNA purification/enrichment, RNA fragmentation, cDNA synthesis, ligation of index adaptors, and amplification, following the KAPA mRNA Hyper Prep Kit Technical Data Sheet, KR1352 – v4.17 (Roche Corporate). Each resulting indexed library was quantified, and its quality accessed by Qubit and Agilent Bioanalyzers; multiple libraries were pooled in equal molarity. The pooled libraries were denatured and neutralized before loading on a NovaSeq 6000 sequencer at 300 pM final concentration for 100b paired-end sequencing (Illumina, Inc.). Approximately 30-40M reads per library were generated. A Phred quality score (Q score) was used to measure the quality of sequencing. More than 90% of the sequencing reads reached Q30 (99.9% base call accuracy). The sequencing data were first assessed using FastQC (Babraham Bioinformatics, Cambridge, UK) for quality control. The sequencing reads were mapped to the mouse genome mm10 using STAR (v2.7.2a) with the following parameter: ‘--outSAMmapqUnique 60’ [18]. Uniquely mapped sequencing reads were assigned to Gencode M22 gene using featureCounts (v1.6.2) [25] with the following parameters: “–p –Q 10 -O”. The genes were kept for further analysis if their read counts > 10 in at least 3 of the samples, followed by the normalization using TMM (trimmed mean of M values) method and subjected to differential expression analysis using edgeR (v3.24.3) [26]. Gene Ontology and KEGG pathway functional enrichment analysis was performed on selected gene sets, e.g., genes undergoing both significant differential expressions and notable changes of open chromatin accessibilities, with the cut-off of false discovery rate (FDR) < 0.05 using DAVID [27]. Canonical pathways from RNAseq data were generated through the use of IPA (QIAGEN Inc., https://www.qiagenbioinformatics.com/products/ingenuity-pathway-analysis) [28]; n=3 samples per condition.

### Assay for Transposase-Accessible Chromatin sequencing (ATACseq)

Cells tested in ATACseq were plated and differentiated to osteoblast/osteocyte-like cells at the same time as those for RNAseq (3 weeks, see above). After differentiation, osteoblast/osteocyte and undifferentiated MSC (control) cells were washed twice in 1X PBS, then dissociated with trypsin (Hyclone) for 5 minutes. Cells were resuspended in ice cold 1X PBS, dead cells were removed, and live cells were processed for nuclei isolation and ATAC sequencing according to published protocols [29]. Briefly, cells were collected in cold PBS and cell membranes were disrupted in cold lysis buffer (10 mM Tris–HCl, pH 7.4, 10 mM NaCl, 3 mM MgCl2 and 0.1% IGEPAL CA-630). The nuclei were pelleted and resuspended in Tn5 enzyme and transposase buffer (Illumina Nextera® DNA library preparation kit, FC-121-1030). The Nextera libraries were amplified using the Nextera® PCR master mix and KAPA biosystems HiFi hotstart readymix successively. AMPure XP beads (Beckman Coulter) were used to purify the transposed DNA and the amplified PCR products. The resulting ATACseq libraries were sequenced on Illumina NovaSeq 6000 and paired-end 50 bp reads were generated. Illumina adapter sequences and low-quality base calls were trimmed off the paired-end reads with Trim Galore v0.4.3. Bowtie2 [30] was used for ATACseq read alignments on the mouse genome (mm10). Duplicated reads were removed using Picard developed by the Broad Institute [https://broadinstitute.github.io/picard/ (Accessed: 2018/02/21; version 2.17.8)]. Low mapping quality reads and mitochondrial reads were discarded in further analysis. Peak calling of mapped ATACseq reads were performed by MACS2 [31] with a Bonferroni adjusted cutoff of p-value less than 0.01. Peaks called from multiple samples were merged, after removing peaks overlapping with ENCODE blacklist regions [32, 33]. Reads locating within merged regions in different samples were counted by pyDNase [34]. The data was filtered using at least 10 cut counts in more than one of the samples, then normalized using TMM (trimmed mean of M values) method and subjected to differential analysis using edgeR (v3.24.3) [26, 35]. Motif enrichment of differential accessibility peaks with a false discovery rate cut-off of 0.05 was performed using Homer [36].

### RNA isolation and qPCR

To isolate total bone RNA, one femur and one tibia per mouse were harvested, the bone marrow flushed using a 27G syringe, and the epiphyses removed, similar to the approach undertaken to prepare the bones prior to collagenase digestion for the scRNAseq studies. The remaining bones (femur and tibia) were harvested and homogenized in 1 ml of TRIzol reagent (Invitrogen) according to the manufacturer’s protocol using a Bullet Blender (Next Advance, Inc.), then further purified using the RNeasy Kit (Qiagen). Mouse *β-actin* was used as an internal control for RT-qPCR. The qPCR primers were purchased as pre-optimized reagents (Applied Biosystems/Life Technologies, Inc.) and the TaqMan One-Step RT-PCR kit was used to perform all reactions. PCR conditions were: 30 minutes 48°C, 10 minutes 95°C, followed by 40 cycles of 15 seconds 95°C and 1 minute 60°C. The data were collected and analyzed by a StepOne Plus system (Applied Biosystems/Life Technologies, Inc.). The expression levels of mRNAs were calculated relative to appropriate controls, and the 2^-ΔΔCT^ method described by Livak was used to analyze the data [37]. The primers used in the study were: Fgf23, Mm00445621_m1; Mmp13, Mm00439491_m1; Tnc, Mm00495662_m1; Gdpd2, Mm00469948_m1; Spp1, Mm00436767_m1; Col1a1, Mm00801666_g1; Bglap, Mm03413826_mH; Cthrc1, Mm01163611_m1; Smpd3, Mm00491359_m1; Dmp1, Mm01208363_m1; Sost, Mm04208528_m1; Pdpn, Mm01348912_g1; Phex, Mm00448119_m1; Ptprz1, Mm00478486_m1; Pdpn(E11), Mm01348912_g1; Actin, Mm02619580_g1; Aldoa, Mm00833172_g1; Adpgk, Mm00511302_m1; Pgam1, Mm02526975_g1; Acadm, Mm01323360_g1; Acp5, Mm00475698_m1; Mmp9, Mm00442991_m1; and Tnfrsf11a, Mm00437132_m1 (Thermo Fisher, Inc).

### Statistical analyses

The most recent updated R software packages with robust affiliated statistics were used to analyze the scRNAseq datasets. The cutoffs used for integration of RNA-seq and ATAC-seq were: peaks withing 10kb upstream of gene, differential peak cutoff FDR < 0.05, differential expression cutoff FDR < 0.05, log_2_FC > 1 (up-regulation) or < -1 (down-regulation). Statistical analyses of the *in vivo* data presented were performed by two-way ANOVA to assess the differences between the same gender in response to adenine diet and to assess the differences between genders and treatments. Significant changes were considered when at *p*<0.05.

## Results

### Mouse osteoblast/osteocyte transcriptomic profiling at single cell resolution

The osteocyte transcriptome at the single cell level solely from long bones remains uncharacterized. To identify cortical bone cells and enrich our cell sample preparation, we used Sclerostin (Sost)-Cre/Ai9 ‘Tomato’ reporter mice at 8 weeks of age to isolate fluorescently-labeled osteoblasts/osteocytes. Consistent with previous characterization [13], SOST-Cre/Ai9 (‘tdTomato’ reporter) mice had detectable basal Cre activation in osteocytes as evidenced by the presence of red fluorescence in mouse tail vertebrae (**Figure 1A**), and were thus harvested in their native state to avoid confounders due to delivery of tamoxifen. Following cortical bone digestion using collagenase/EDTA, the cells were sorted, and the Ai9/tdTomato^+^ fraction represented approximately 1.5% of total cells (**Figure 1B**). As determined by qPCR, the Ai9/tdTomato^+^ cells were highly enriched for osteocyte marker mRNAs *Fgf23* (40-fold), *Phex* (102-fold), *Dmp1* (172-fold), and *Pdpn* (12-fold), compared to the Ai9/tdTomato^-^ population (**Figure 1B**).

**Figure 1.**
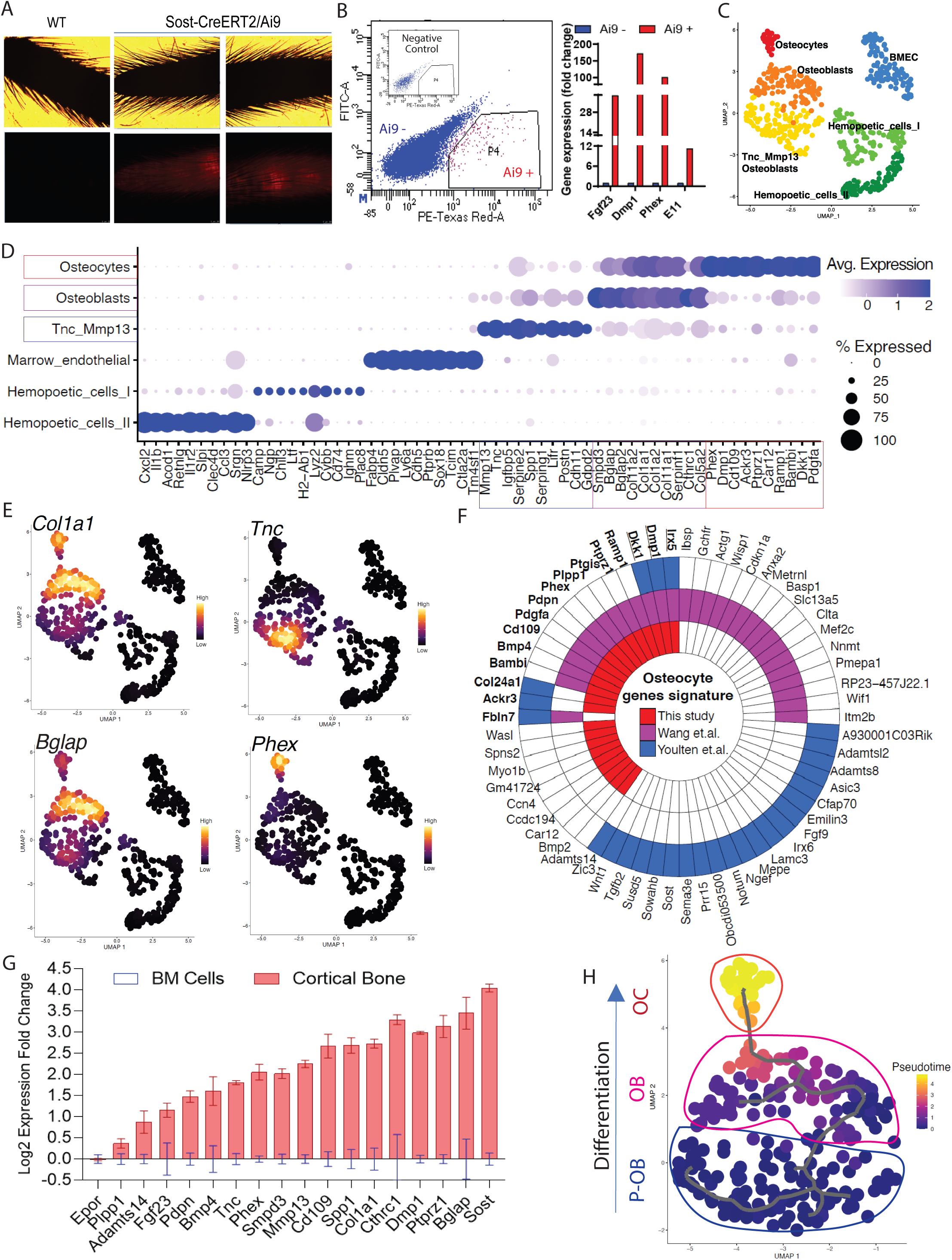
Single-cell transcriptomic profiling of bone cells identified osteolineage heterogeneity. **A-B.** Long bones (femur, tibia, and humeri) were subjected to serial collagenase/EDTA digestions (see Methods), and cells from fractions 4-9 were collected for FACS sorting to isolate tdTomato-positive cells, followed by single-cell RNAseq library construction. Analysis by qPCR confirmed high mRNA expression of osteocyte genes *Fgf23*, *Dmp1*, *Phex*, and *Pdpn* in Ai9 positive cells *versus* Ai9 negative cells. **C.** Bioinformatic analysis was performed to cluster the cells by UMAP. Each dot represents a single cell, and cells sharing the same color code indicate discrete populations of transcriptionally similar cells. **D.** Defining cortical bone cell types. Tnc/Mmp13 osteoblast cells were distinctly marked by *Tnc* and *Mmp13*. Osteoblast cells showed high expression of *Smpd3* and *Bglap*. Osteocytes had the highest expression of *Dmp1* and *Phex*. **E.** Expression density plots indicated cells with high transcription of *Col1a1*, *Tnc*, *Bglap*, and *Phex*. **F.** The polar figure highlights the osteocyte markers isolated from our dataset (long bone cell scRNAseq), Wang et al. (long bone and calvaria cell scRNAseq; [10]) and Youlten et al (long bone bulk RNAseq; [9])**. G.** Canonical and non-canonical osteolineage genes were validated as being highly expressed in cortical bone when compared to the expression detected in bone marrow. **H.** Pseudotime analysis revealed Tnc/Mmp13 Osteoblasts as an osteoblast precursor cell (P-OB), and shows their differentiation to osteoblasts (OB) and then osteocytes (OC).

To identify the transcriptome of the Ai9/tdTomato^+^ population, scRNAseq was performed on this isolated cell population. Approximately 1,200 cells were successfully recovered after sequencing with an estimation of 130,000 reads per cell and 97% valid barcodes. Further stringent bioinformatic analyses were performed to exclude low-quality cells; for downstream analysis, cells that had more than 3500 genes were excluded to reduce doublet nuclei, and less than 15% of mitochondrial genes to exclude potentially dead cells (**Figure S1A**). To aggregate cells based on their similarities, principal component analysis (PCA) was performed on highly variable genes and the result was used as input for clustering using the Louvain algorithm with multilevel refinement and Uniform Manifold Approximation and Projection (UMAP) for dimension reduction (**Figure 1C**). To identify gene specific markers in each cluster, we used the function FindMarkersAll from the MAST method algorithm. The quality of clustering was assessed by dot plot of the most significant marker genes obtained from an individual cluster (**Figure 1D**).

The transcriptomic profiling of Ai9/tdTomato^+^ cortical bone cells identified a total of six cell populations, including two osteoblast clusters and one osteocyte cluster (**Figures 1C and 1D**). The sequenced cells were grouped into three partitions: hematopoietic cells that are Cd45 (Ptprc) positive, endothelial cells that highly express Cdh5, and osteolineage that express the pan mesenchymal lineage cell marker Pdgfra, which was found to be present in pre-osteoblasts, osteoblasts and osteocytes (**Figure S1B-D**). Marker analysis revealed that within the osteoblast population, one cell type was characterized by higher expression of *Smpd3*, *Bglap, Col1a1*, and *Col11a1* mRNAs. The second, referred to as “Tnc/Mmp13 osteoblasts,” was defined by high expression of *Tnc*, *Mmp13, Serping1 and Spp1*, which was consistent with previous subpopulation analyses [10]. Osteocytes were associated with a cluster showing the highest expression of *Phex* and *Dmp1*. Other genes that defined osteocytes were *Cd109*, *Dkk1*, and *Ptprz1* (**Figure 1D,** and osteolineage markers listed in **Supplementary Table 1**). Our analyses also identified two distinct populations of hematopoietic cells. The first population annotated “Hematopoietic I” showed high expression of *Cybb*, *Chil3*, and *Ltf* whereas the “Hematopoietic II” cells were characterized by expression of *Clec4d*, *Il1r2*, and *Cxcl2*. Bone marrow derived endothelial cells (‘BMEC’) were also identified and distinctly defined by *Cdh5*, *Plvap*, and *Ptprb* gene transcripts (**Figure 1D**). As expected, *Col1a1* was identified only in osteolineage cells (**Figure 1E**), underscoring the ability to identify transcripts that may be uniquely associated with ossification.

Our UMAP analysis confirmed that the Tnc/Mmp13 osteoblast population with high Tnc, Mmp13 and Spp1 expression (**Figure 1E and Figure S1E-F**) clustered farther from osteocytes when compared to the dimensional separation between the defined more mature osteoblasts (high *Bglap* expression (**Figure 1E**)) and osteocytes (high *Phex* expression (**Figure 1E**)). These data suggest that the Tnc-Mmp13 osteoblasts likely represented a precursor osteoblast population. Importantly, we isolated 25-highly and preferentially expressed genes in osteocytes, with some overlap of genes in a proposed osteocyte transcriptome [9]. The polar plot comparison of recently reported osteocyte transcriptome highlights the genes in common such as *Dkk1*, *Dmp1*, *Irx5, Col24a1, Ackr3* from our dataset (scRNAseq of long bone), Wang and colleagues (scRNAseq on long bone and calvaria) [10] and Youlten and colleagues (bulk RNAseq on long bone) [9] (**Figure 1F**).

To further test whether the genes identified as highly expressed in osteolineage cells in our scRNAseq analyses were not derived from bone marrow cells, we compared the mRNA expression of these candidate genes in mouse cortical long bone (osteoblast/osteocyte-enriched fraction) *versus* bone marrow using qPCR. In this regard, *Mmp13*, *Cthrc1*, *Smpd3*, *Ptprz1*, and *Cd109* were tested, and these genes were highly expressed in cortical bone suggesting that they were preferentially expressed in osteolineage cells (**Figure 1G**). The erythropoietin receptor (*Epor)* mRNA, tested as a marrow positive control, was not increased in cortical bone when compared to bone marrow expression levels, whereas defined osteocyte genes *Phex, Pdpn, Dmp1, Fgf23,* and *Sost* were all increased in cortical bone samples (**Figure 1G**). To determine the cellular differentiation patterns that govern the maturation of osteoblasts to osteocytes, we performed pseudotime analysis. This algorithm was first performed on the partition of cells based on the UMAP (**Figure 1C**) in three main compartments (p1, osteolineage; p2, hematopoietic cells; and p3, marrow endothelial cells) as indicated in **Figure S1G**. After the initial analyses, Monocle3 simultaneously performed pseudotime analysis over the individual partitions. Trajectory analyses from the partition p1, an osteogenic lineage (**Figure 1H**), hematopoietic cells (**Figure S1H**), and endothelial cells (**Figure S1I**) were next generated. Interestingly, a global cellular interaction pattern was conserved with almost the same UMAP projection as for all cell populations (as in **Figure 1C**). These analyses confirmed that the osteocyte cluster of cells identified from our dataset derived from the osteoblast subset, with the Tnc/Mmp13 osteoblast as a likely precursor population (**Figure 1H**).

### scRNAseq identifies pathways involved in the osteoblast to osteocyte transition

To identify the molecular pathways involved in the transition from osteoblasts to osteocytes at single cell resolution, we analyzed differentially expressed osteoblast/osteocyte genes using Ingenuity Pathway Analysis (IPA). Further, a comparison analysis was performed to identify signaling pathways in osteoblasts *versus* osteocytes. In this regard, we found an enrichment of “*GP6 Signaling*”, a major signaling receptor for collagen in osteoblasts, which was almost completely shut down in osteocytes (**Figure 2A**). In contrast, the “*Axonal guidance*”, the “*Differentiation via BMP receptors*”, “*TGFβ signaling*” and the “*Iron homeostasis signaling*” pathways that were not highly enriched in osteoblasts were significantly increased in osteocytes (**Figure 2A**). Consistent with osteocyte morphology, among the genes predicted to drive “*Axonal guidance*” were *Adamts14*, *Bmp2*, *Bmp4*, and *Wasl*. Furthermore, IPA analysis predicted EGFR among the top upstream regulators of osteoblastic pathways whereas FGF2 was predicted to be a primary upstream regulator associated with the osteocyte maturation pathway (**Figure S2A**). Additionally, our in-silico analysis identified an enriched cell activation in osteoblasts, associated with genes such as *Col11a1*, *Col11a2*, and *Col1a1* which were significantly attenuated in osteocytes (**Figure 2A**). Overall, these data support that osteoblasts and osteocytes can be defined by unique pathways associated with their individual homeostatic functions.

**Figure 2.**
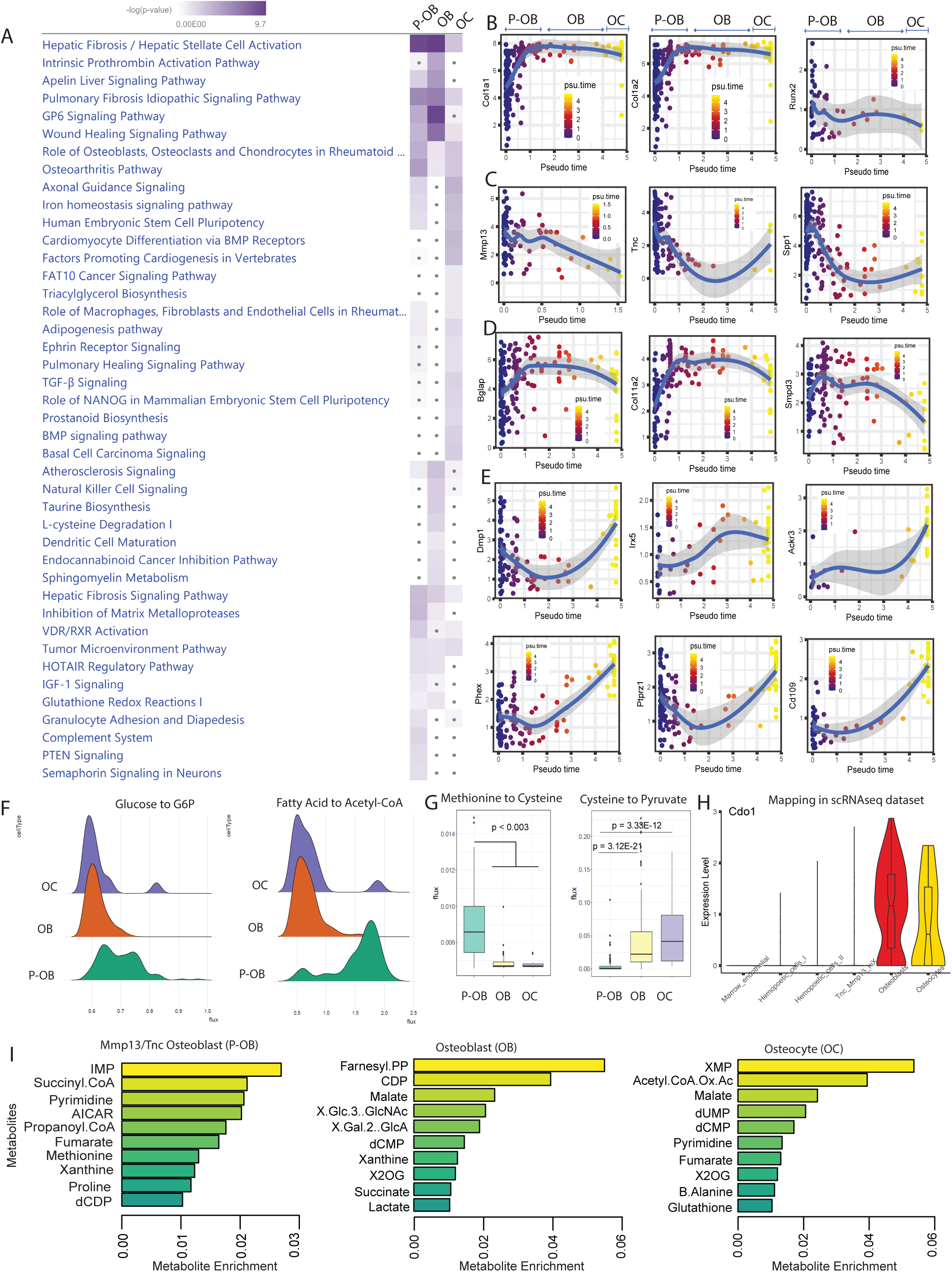
Differential signaling pathways in osteoblasts versus osteocytes. **A.** Ingenuity Pathway Analysis (IPA) software was used to predict the statistically significant canonical pathways upregulated in Tnc/Mmp13 osteoblasts (P-OB), osteoblasts (OB) and osteocytes (OC) from the scRNAseq data based upon a calculated probability score of ≥2 and a p-value of <0.05. Further, a comparative analysis between osteolineage cells was performed to generate the heatmap. The increase of blue intensity reflects the more significant pathways within cell types. The gray dots indicate non-significant pathways. **B-E.** Monocle analysis shows pseudotime trajectory mapping of osteolineage gene sets *Col1a1, Col1a2, Runx2;* pre-osteoblast gene sets *Mmp13, Tnc, Spp1;* osteoblast gene sets *Bglap, Col11a2, Smpd3;* and osteocyte gene sets *Dmp1, Irx5, Ackr3, Phex, Ptprz1,* and *Cd109*. **F.** The computational method scFEA was used to infer cell-wise fluxome from the scRNAseq data to predict the metabolic profiling as well as the differential metabolite conversion rate in cells that correspond to a metabolic flux value. Ridgeline plots indicate the distribution values of metabolic flux in Tnc/Mmp13 osteoblasts (P-OB), osteoblasts (OB), and osteocytes (OC). Each ridgeline represents the flux between two metabolites, shown on the x-axis, for different osteolineage cells, shown on the y-axis. **G.** Violin plot displaying *Cdo1* expression in scRNAseq dataset. **I.** The boxplots indicate the predictive value of the conversion of methionine to cysteine and to pyruvate in P-OB, OB, and OC. **H.** The bar plots show the top 10 different metabolites enriched in P-OB, OB, and OC.

To predict the transcriptional profiles of osteocyte genes across the osteolineage, we used pseudotime analysis to track the computational-derived kinetics of gene expression during differentiation. We found that “pan-osteolineage” markers *Col1a1*, *Col1a2* and *Runx2* showed relatively stable expression during osteoblast to osteocyte differentiation (**Figure 2B**). In contrast, *Mmp13, Tnc,* and *Spp1* pseudotemporal progression were predicted to decrease during differentiation from Tnc/Mmp13 osteoblasts to osteocytes (**Figure 2C**). Further, *Bglap, Col11a2,* and *Smpd3* expression increased from the Tnc/Mmp13 precursor osteoblasts to osteoblasts, but decreased when the osteoblast population transitioned to osteocytes (**Figure 2D**). In osteocytes, the kinetic profiling of *Dmp1, Irx5, Ackr3, Phex, Ptprz1,* and *Cd109* showed progressive positive regulation during differentiation (**Figure 2E, Figure S2B,** and **Supplementary Table 1**). These data confirmed the dynamic differentiation of canonical genes during the transcriptional reprogramming required for osteoblast transition to osteocytes, as well as identified new genes associated with the cortical bone osteolineage.

To distinguish the metabolic heterogeneity between osteoblasts and osteocytes, we applied the recently developed computational method single-cell flux estimation analysis (scFEA) [23] that uses a systematically reconstructed human metabolic map to infer the cell-wise cascade from the transcriptome to metabolome. This approach applies multilayer neural networks to capitulate the nonlinear dependency between enzymatic gene expression and reaction rate fluxome from scRNAseq datasets. By reconducting the cell clustering analysis using the original UMAP labeling (**Figure 1C**), scFEA identified high metabolic reactions including glycolytic, TCA, and fatty acid metabolic reactions in Tnc/Mmp13 osteoblasts (**Figure S2C-S2E**) consistent with an overall increased cell activation (**Figure 2A**). The most prominent change of metabolic route predicted was the conversion of Fatty acid to Acetyl co-A during the transition from pre-osteoblasts to osteoblasts/osteocytes with two distinct cell populations (**Figure 2F**). The gene *Acadm* (Acyl-CoA Dehydrogenase Medium Chain) was the top predicted gene to drive this process. Whereas the conversion of serine to cysteine decreased with osteoblast differentiation, the transformation of cysteine to pyruvate increased when Tnc/Mmp13 osteoblasts transitioned to osteoblasts/osteocytes, a potential alternative metabolic route to supply cells with pyruvate (**Figure 2G**). This cysteine metabolism change predicted by scFEA correlated with differential regulation of *Cdo1* (Cysteine Dioxygenase Type 1), a gene that was highly expressed in osteoblasts (**Figure 2H**). Based on the scRNAseq dataset, the top predicted metabolites in osteolineage cells were the inosine monophosphate (IMP), succinyl coenzyme A, and pyrimidine in pre-osteoblasts; the farnesyl pyrophosphate (FPP), cytidine-5’-diphosphate (CDP), and malate in osteoblasts; whereas the xanthosine monophosphate (XMP), acetyl CoA oxaloacetate, and malate were enriched in osteocytes (**Figure 2I**). Collectively, the transcriptional profile across cortical bone cell types predicted differential metabolic states between the osteoblast cell population subsets and osteocytes.

### Identification of genes during staged osteolineage differentiation

To validate the osteoblast/osteocyte genes identified from our *in vivo* scRNAseq experiments, the mRNA expression of these genes was tested in mesenchymal progenitor cell (‘MPC2’) cells *in vitro*. MPC2 cells were recently characterized as harboring the ability to derive osteoblast- and osteocyte-like cells when cultured in osteogenic media [24]. After 1-4 weeks of culture, the differentiated cells can be stained by alizarin red reflecting the ability of the mature cells to mineralize [24]. Our *in vitro* molecular analyses demonstrated temporal changes of canonical and non-canonical osteolineage genes. *Col1a1* mRNA increased during cell differentiation with the highest level observed at 4 weeks. *Runx2* gene expression reached an early and maximal expression 1-week after differentiation and remained stable until 4 weeks. The expression of Tnc significantly increased 1-week after differentiation, although *Cthrc1,* identified as an enriched osteoblast gene based upon our scRNAseq dataset began increasing with MPC2 cell differentiation at 3 weeks (maximal time point) before being significantly downregulated at 4 weeks (**Figure 3A**). Osteocyte genes, including *Pdpn*, *Phex*, *Sost* and *Ptprz1,* had high expression at 3 and 4 weeks of osteocyte differentiation, confirming our finding of *Ptprz1* as a likely late osteoblast/osteocyte gene. Further, genes associated with predictive glycolytic and β-oxidation pathways identified by scFEA from the *in vivo* studies (**Figure 2F**) were assessed during *in vitro* MPC2-osteoblast differentiation. We found that glycolytic genes increased during osteoblast differentiation including *Adpgk* (ADP-Dependent Glucokinase)*, Aldoa* (Aldolase A), and *Pgam1* (Phosphoglycerate Mutase 1) reaching ∼3-fold increases at 3 at 4 weeks. In contrast, *Acadm* (Acyl-CoA Dehydrogenase Medium Chain), a gene associated with the conversion of fatty acid to acetyl co-A remained stable during the first three weeks of differentiation before being downregulated at 4 weeks (**Figure 3A**), consistent with the predicted osteocyte metabolic transition *in vivo* (**Figure 2F**).

**Figure 3.**
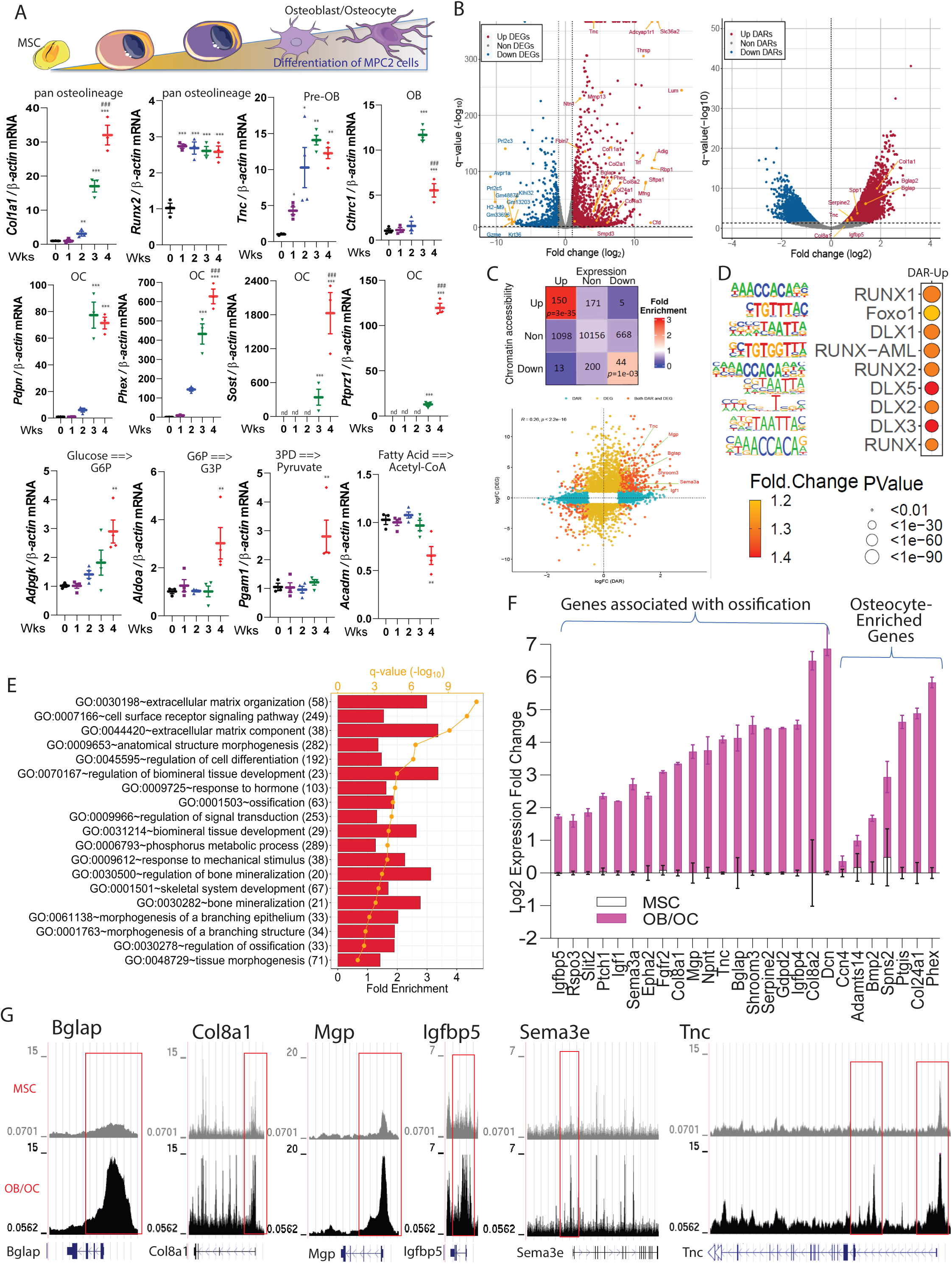
Molecular markers associated with osteoblast/osteocyte differentiation. **A-top.** Schematic representing cell differentiation from MSCs to osteoblasts/osteocytes. **A-lower.** *Col1a1*, *Runx2, Tnc, Cthrc1, Pdpn, Phex, Sost, Ptprz1, Adpgk, Pgam1, Aldoa,* and *Acadm* mRNAs were analyzed in MPC2 cells during osteogenic differentiation over the course of 1-4 weeks and normalized to β-actin. Data are represented as mean +/- standard deviation. **p<0.01, and ***p<0.001 compared to Control (0 Wks). ^###^p<0.001 compared to 3 Wks. nd = not detected. **B.** Volcano plots derived from the bulk RNAseq (left) and ATACseq datasets (right) show the significant alterations of gene expression and chromatin accessibility detected in differentiated *versus* undifferentiated cells. The x-axis shows the fold-change (in log_2_ scale). The y-axis represents the significance. **C-top.** The 3 Χ 3 matrix heatmap highlights the number of genes with differentially accessible regions (DARs), or differentially expressed genes (DEGs), or both DARs and DEGs when compared differentiated *versus* undifferentiated cells. **C-lower.** Integration of ATACseq and RNAseq data exhibited a higher correlation between genome-wide chromatin accessibility changes (x-axis) and gene expression alterations (y-axis). **D.** Selected motifs of more open regions of the genome in differentiated cells *versus* undifferentiated cells using the ATACseq. **E.** Functional enrichment analysis using the Database for Annotation, Visualization and Integrated Discovery (DAVID) on upregulated DEGs with more open DARs identified biological pathways that were induced during osteogenic differentiation (numbers in parenthesis are gene counts). **F.** Representative mRNA expression of upregulated genes in differentiated versus undifferentiated cells enriched in cell morphogenesis and ossification. All the genes shown were significantly upregulated (false discovery rate, FDR < 0.05). The osteocyte genes identified from the scRNAseq dataset in Figure 1J were confirmed to be significantly upregulated in osteoblasts/osteocytes cells *versus* control undifferentiated MPC2 cells. **G.** Representative ATACseq peaks of undifferentiated cells (top track in gray) compared to differentiated cells (osteoblasts/osteocytes bottom track in black) are shown.

To identify the molecular events associated with osteolineage differentiation, a comprehensive genomic and transcriptomic analyses using RNAseq and ATACseq was employed using MPC2 cells [38]. Based on the kinetic profile performed in **Figure 3A**, a time point of 3 weeks of MSC to osteocyte-like cell differentiation was chosen for RNAseq and ATACseq to capture differentially regulated genes in osteoblasts and osteocytes. Our unbiased approach identified several genes with increased chromatin accessibility and/or gene expression during MPC2 differentiation including osteocalcin (*Bglap*), osteopontin *(Spp1), Col11a1,* and *Col11a2* (**Figure 3B** and **Figure S3A-F**). By performing integrative and correlative analyses of the ATACseq and RNAseq, we confirmed our observed changes in mRNA expression and chromatin accessibility in the ATAC-seq dataset. Using this approach, 150 genes were identified with significantly more open chromatin status and higher gene expression levels after differentiation (**Figure 3C**), whereas 44 genes were downregulated with notably less DNA accessibility (**Figure 3C**). The gene sets with both upregulated differential chromatin accessible regions (DAR) and differential expressed genes (DGE), as well as downregulated DAR and DE showed a significantly high fold-enrichment compared to the random selections. However, all other combinations did not demonstrate significance *versus* the random selections (**Figure 3C**). Further, a significant Jaccard index score was noted when gene expression and chromatin accessibility increased or decreased simultaneously (FDR < 0.05, see **Supplementary Table 2**), suggesting that the chromatin changes reflected by DARs, tended to be positively correlated with the DEGs. HOMER transcription factor motif analysis of the ATACseq data detected an enrichment of several motifs that are known to influence bone differentiation including an increased enrichment of Runx1, Runx2, Foxo1, and DLX1/5/2 (**Figure 3D**).

The upregulated genes identified from differential expression analysis associated with changes in chromatin accessibility (**Figure 3D**) were then subdivided by gene ontology (GO). Among the GO terms that were increased during osteoblast/osteocyte differentiation were functions related to bone mineralization, as well as cellular morphology changes including membrane branching (**Figure 3E**). Genes that reflected pathways associated with “Cell Morphology” and “Ossification” are shown in **Figure 3F**. Importantly, osteocyte genes identified from the scRNAseq analysis including *Ccn4, Adamts14, Spns2,* and *Bmp2* were significantly upregulated in osteocyte-like MPC2 cells when compared with parent undifferentiated MSC cells (**Figure 3F**). *Ptgis* identified in our dataset and the Wang, et al dataset [10] was also significantly increased with differentiation. *Col24a1* found in our dataset and the Youlten, et al dataset [9] was also increased (**Figure 3F**). These mRNA expression levels detected during the cell transitions were robustly and positively associated with their corresponding genomic changes (**Figure 3G**). For example, the DNA promoter regions at *Bglap*, *Col8a1*, *Mgp*, *Igfbp5*, *Sema3a* and *Tnc* genes were more accessible with cell maturation (**Figure 3G**).

Next, we tested the chromatin accessibility of a set of osteolineage genes identified in the scRNAseq. For instance, among the Tnc/Mmp13 osteoblast signature genes, we detected increased chromatin accessibility at the *Mmp13*, *Serpine2,* and *Lifr* promoter regions in differentiated cells versus MSC control cells (**Figure S4A-C**). We also observed high chromatin accessibility at the promoters of *Serpinf1* and *Cthrc1* (**Figure S4D-E**), that were defined as expressed by fully differentiated osteoblasts based upon our *in vivo* scRNAseq data (see **Figure 1D**). For osteocyte genes, increased chromatin accessibility for the promoter regions of *Ackr3, Cd109, Ptgis, Spns2,* and *Bmp2* was observed (**Figure S4F-J**). Taken together, these results identified genomic regions of interest associated with osteoblast differentiation that may play critical roles in cellular transcriptional reprogramming.

### Impairment of osteoblast/osteocyte genes prior to cortical bone deterioration in CKD

We next assessed the dynamic transcriptional profile of the identified osteoblast and osteocyte genes within cortical bone prior to the development of cortical porosity in CKD, a severe outcome that leads to bone fracture. We used a mouse model in which CKD is induced by providing an adenine-containing diet, which causes a progressive loss of kidney function [39]. This model reflects the phenotype observed in humans, characterized by progressive tubular atrophy, immune cell infiltration, and fibrosis [39], and has been extensively characterized as a reliable model to investigate CKD-mineral and bone disorder [15]. In comparison to healthy controls, previous studies showed that mice fed a 0.2% adenine diet exhibited biochemistries typical to CKD such as hyperphosphatemia, hypocalcemia, secondary hyperparathyroidism, increased blood urea nitrogen (BUN), and FGF23 induction, which are enhanced with the duration of treatment, as well as associated with bone porosity during chronic adenine administration [15].

With a focus on testing osteoblast/osteocyte gene regulation in cortical bone at a timepoint prior to major ultrastructural changes, cohorts of C57BL/6 mice were placed on control diet or a 0.2% adenine-containing diet for 2 or 4 weeks. At 2 weeks, serum phosphate, alkaline phosphatase, and calcium were not significantly changed compared to controls (**Table 1**). Blood urea nitrogen (BUN) was monitored for declines in renal function, and as expected, the mice receiving the adenine diet had significantly increased BUN (**Table 1**). The cortical porosity and bone volume remained unchanged at 2 or 4 weeks in male or female CKD mice (**Figure 4C**). However, plasma bioactive iFGF23 concentrations were markedly elevated in the mice with CKD (**Figure 4D**).

**Figure 4.**
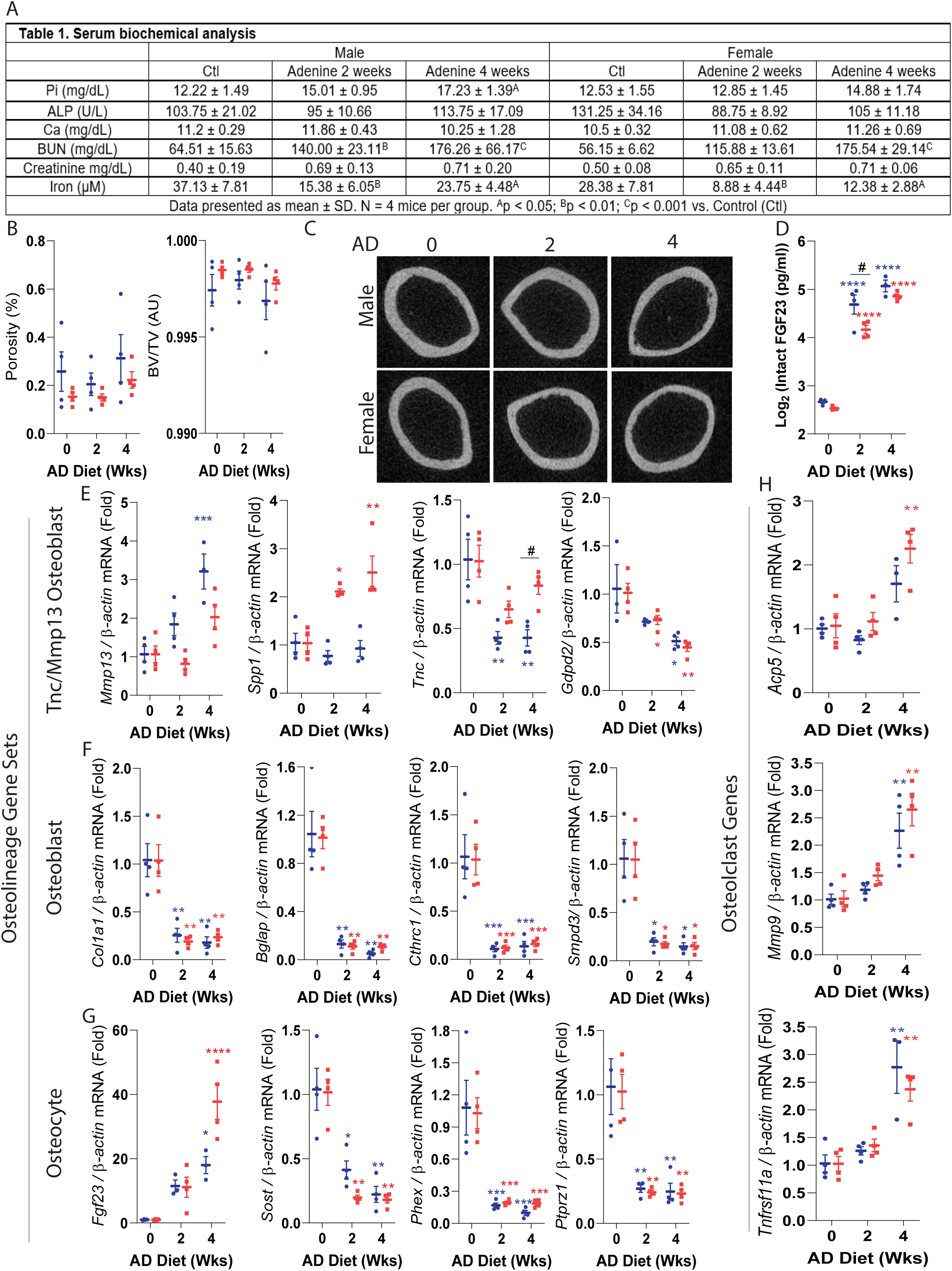
Precocious misregulation of osteoblast/osteocyte genes in CKD. Eight-week old wild type C57BL/6 male (shown in blue) and female (represented in red) mice were fed an adenine diet (AD) to induce CKD for 2, or 4 weeks. Mice fed the casein diet for 2 weeks were used as controls. **A.** Key serum biochemical analyses: phosphate, alkaline phosphatase, calcium, blood urea nitrogen, creatinine, and iron are shown. **B.** Cortical porosity (left) and trabecular bone volume (right) were measured using micro computed tomography (μCT). **C.** Representative images of cortical bone μCT. **D.** Circulating FGF23 was assessed by ELISA. **E-H.** Real-time qPCR was used to measure the mRNA expression of Tnc/Mmp13 osteoblast (pre-osteoblast), osteoblast and osteocyte gene sets, as well as osteoclast genes. Data are shown as fold change (2-ΔΔCt) relative to the housekeeping gene *β-Actin* and normalized to the experimental control (‘AD0’). The ‘AD 0’ mice were fed with a control diet (Casein) and sacrificed at 2 weeks; ‘AD 2’ mice were fed with a CKD diet (adenine) and sacrificed at 2 weeks; and the ‘AD 4’ were mice fed with the adenine diet and sacrificed at 4 weeks. Data are represented as mean +/- standard deviation. *p<0.05, **p<0.01, ***p<0.001, ****p<0.001 compared to Control.

We next determined the effects of CKD on highly expressed osteoblast and osteocyte genes in cortical bone samples. During early CKD, *Mmp13*, and *Spp1* identified in the osteoblast precursor population, remained stable or increased with CKD (**Figure 4E**). In contrast, other precursor osteoblast genes such as *Tnc,* and *Gdpd2,* were decreased during CKD progression. The expression of the osteolineage gene *Col1a1*, as well as osteoblast-specific genes *Bglap, Cthrc1, Smpd3* and osteoblast/osteocyte marker *Dmp1*, were strongly impaired in the mice with CKD (**Figure 4G**). Increased circulating FGF23 was associated with significant upregulation of cortical bone *Fgf23* mRNA at 4 weeks (**Figure 4G**). In a similar pattern to that of osteoblast genes, osteocyte genes including *Sost*, *Phex,* and *Ptprz1* were dramatically downregulated early in CKD (**Figure 4G**). To test whether the change of osteocyte genes in early CKD resulted from an overall decrease of cell numbers, the osteocytes were counted in cortical and trabecular bone using histological analysis. Osteocytes were found within lacunae, and their cell numbers were not different between controls and mice with CKD (**Figure S5A-B**), suggesting that the observed changes in gene expression in cortical bone were not due to the decrease of overall cell numbers but rather a profound change at cellular level. Of note, empty lacunae were not observed. The osteocyte numbers in trabecular bones were also unchanged, however the number of osteoblasts in trabecular bone was increased in the mice with CKD (**Figure S5A-B**).

To assess whether changes in bone forming genes at CKD onset occur concurrently with changes in osteoclast markers, we measured gene expression of *Acp5* (Tartrate-resistant acid phosphatase), *Mmp9* (Matrix metallopeptidase 9), and *Tnfrsf11a* (Rank) by qPCR. At 2 weeks, a time point where serum phosphate remained unchanged and the cortical bone unaltered as tested by µCT, the mRNA expression of these osteoclast markers was not different from controls (**Figure 4H**). However, by 4 weeks, these genes were upregulated, consistent with an onset of metabolic bone disease. These findings suggest that at the early stage of the disease, the misregulation of bone-forming osteolineage genes may be independent from loss of cortical bone or increased osteoclast function.

In sum, this work demonstrated that scRNAseq can identify sub-populations of osteoblasts, and together with metabolic profiling, predict differentiation to mature osteoblasts and osteocytes. In addition, we identified genes not previously associated with cortical bone expression, and confirmed their presence in osteoblast/osteocytes via corresponding changes in genomic accessibility and expression using *in vitro* differentiation studies. These genes were then shown to be misregulated in a mouse model of CKD. Our collective findings support that the molecular changes observed in cortical bone associated with the severe musculoskeletal phenotypes in CKD may occur more rapidly than recognized, and can now be pinpointed to specific cell types within the osteolineage.

## Discussion

There is an urgent need to understand bone-forming cell heterogeneity as well as isolate key gene networks that impact skeletal homeostasis. This is especially apparent in diseases such as CKD, where due to the progressive nature and specific calcium- and phosphate-related endocrine disturbances, there are currently very little treatment options [40, 41]. Our cortical bone single cell RNAseq analysis identified two osteoblast populations as well as a distinct osteocyte cluster. The genes demarcating each cell type were confirmed as highly enriched in cortical bone as well as being upregulated during osteoblast differentiation *in vitro*. Interestingly and unexpectedly, at early-stage CKD onset in mice, there was a dramatic misregulation of gene expression in cortical bone cell types prior to major bone ultrastructural changes.

Recent studies using bulk RNA sequencing developed an osteocyte-defining transcriptome which was comprised of 28 genes [9], and we observed partial overlap of genes within the transcriptomes reported by Youlten et al [9], as well as from analysis we performed using the publicly available scRNAseq dataset of Wang et al [10]. Our scRNAseq analysis also identified additional genes which were highly enriched in osteocytes (see **Figure 1F**). It is likely that some of the observed differences in osteocyte gene profiling between our studies and those of others are due to the fact that we exclusively used long bones for cell isolation whereas the previous studies sequenced a mix of long bone and calvariae, or used bulk total cortical bone RNA as a starting material [9, 10]. Future studies will be needed to further sub-set these gene profiles and expand cell-specific targets as more cortical bone datasets become publicly available for multi-study integration. This is especially the case for osteolineage genes in light of the idea that these cells are derived from precursors that have distinct and overlapping gene expression.

Our bone scRNAseq analyses identified a population of Tnc/Mmp13 mRNA-containing osteoblasts as potential precursors of a mature osteoblast population of cells. Mmp13, the most highly expressed gene in this cell type has been described as a direct target of osteoblast-specific transcription factor osterix (Sp7) in osteoblasts [42] and known to be activated by Runx2, the master transcription factor for osteoblastogenesis [43]. A recent study defined the molecular mechanisms driven by Sp7 to control osteocyte dendrite formation [10]. Whether directly targeting Mmp13 could influence osteoblast differentiation and osteocyte maturation remains to be investigated. Further analyses integrating our bone cell data sets with other published scRNAseq data by including stem cells and osteoprogenitors may complement the work described herein to predict a full osteogenic trajectory, and to precisely position the Tnc-Mmp13 osteoblast population in the osteogenic commitment lineage. Although we identified osteolineage genes in common with previous studies [9, 10] in addition to unique markers, we cannot exclude that using a system of basal state SOST-Cre/Ai9 and cell sorting (with no tamoxifen induction) identified a specific set of cortical bone cells. It will be important to continue to characterize additional bone cell populations identified through multiple approaches. As new tools such as mouse reporter genes for studying osteoblasts/osteocytes continue to emerge in concert with evolving approaches for testing the single cell transcriptome and genomic accessibility with more sensitivity, it will be important to continue to refine bone cell classification to fully understand osteolineage heterogeneity.

Our work in modeled CKD identified a general misregulation of mature osteoblast and osteocyte genes prior to major bone structural changes. This analysis detected a downregulation of multiple osteoblast and osteocyte mRNAs except for bone *Fgf23* that increased in CKD, or remained stable or were moderately induced, such as *Mmp13* and *Spp1*. The pre-osteoblast gene sets tested in CKD suggested a step-wise decrease in progenitor alterations. Further, our histological analyses found that the number of osteocytes in cortical and trabecular bone are unchanged at the early stage of modeled CKD, suggesting that diminishing cell numbers with disease progression was not responsible for the altered gene expression (**Figure S5A-B**). These fundings are consistent with previous studies that quantified osteocytes in healthy controls, pre-dialysis CKD patients, and pediatric dialysis patients. This report found that the osteocyte cell numbers in bone biopsies did not differ between CKD patients when compared to the healthy group [44]. In support of our findings, it is possible that the maturation mechanisms of osteocytes may instead be affected [44]. Thus, the genes tested herein may serve as early onset markers for CKD bone disease and set the stage for studies that could determine their function during osteolineage differentiation in CKD. These studies also imply that targeting one bone cell type in CKD bone disease will likely not be fully effective since damage may be done in early precursor cells, with these cells potentially carrying these gene expression changes through differentiation where they may impact mature cells. We also found a decrease in cortical bone Sclerostin (*Sost*) expression at an early stage of CKD in mice (**Figure 4G**). In a paracrine signaling manner, rapid decreases of osteocyte genes *Sost* and likely *Dkk1* may represent an adaptive mechanism to maintain effective Wnt signaling as an attempt to delay the initial development of renal osteodystrophy. Whether the decrease in cortical bone *Sost* stimulates trabecular osteoblast differentiation in a paracrine manner to attempt to sustain bone formation at CKD onset remains to be determined.

It was demonstrated in previous studies that mice fed an adenine-containing diet over a chronic 10-week period developed deficits in cortical bone properties associated with an increase of osteoclast surface and a higher bone formation rate [45]. Another adenine study conducted over a 56-day protocol found that the bone resorption marker carboxy-terminal collagen crosslinks (CTX) was decreased and the bone formation marker the N-terminal propeptide of type I procollagen (P1NP) had a trend towards increasing (p=0.06) [46]. These chronic CKD models suggested a high bone turnover phenotype when mice were fed with an adenine diet over a long-term period. In our studies, mice exposed to adenine diet for only 2 weeks did not exhibit cortical porosity, and osteoclast gene markers were unchanged at 2 weeks of dietary treatment although osteoblast and osteocyte markers assessed in cortical bone were already downregulated (**Figure 4F-H**). These results suggest that misregulation of genes within the osteolineage may precede the up-regulation osteoclast genes.

The skeleton is an important regulator of systemic glucose homeostasis, with osteocalcin and insulin thought to represent prime mediators of the interplay between bone and energy metabolism [47, 48]. Using the recently developed computational tool single-cell flux estimation analysis (scFEA) [23] that calculates the cell metabolomic state in scRNAseq datasets, our data predicted *in vivo* metabolic transitions during osteoblast differentiation. At the single cell level, our data supported high glycolysis and glutamine metabolic states in osteoblast precursors versus more mature osteoblasts and osteocytes (**Figure 2F** and **Figure S2G**). The metabolic flux analysis suggested that metabolomic transition between pre-osteoblasts to osteoblasts was more pronounced than between osteoblasts to osteocytes. This was correlated with the extent of differentially expressed genes between osteolineage cells. Indeed, 198 genes were differentially expressed when comparing pre-osteoblasts and osteoblasts *versus* 77 genes between osteoblasts and osteocytes (logFC>0.5 and p<0.05; **Figure S2G**). The conversion of arginine to ornithine was similar across all osteolineage cells (**Figure S2H**), and pathways including the conversion of citrulline/aspartate to arginosuccinate or acetyl glucosamine to hyaluronic acid were also predicted to be conserved regardless of cell differentiation status or cell type (**Figure S2H**). In our analyses, we provided validation in a distinct cell system for the regulation of key genes associated with these metabolic flux predictive data. However, it will be important to build upon these findings to provide genetic evidence that specific osteoblast precursor genes assorted with metabolism may be critical in supporting osteoblast differentiation, as well as influencing global energy metabolism in normal states and in skeletal disease.

Although this study provided an extensive gene profile analysis of osteolineage cells, some established osteocyte genes such as *Sost* and *Fgf23* were not detected under baseline conditions. It is likely that current single cell sequencing technologies using the short coverage length along each mRNA from the 3’UTR end (which is based on 28 bp of cell barcode and UMI sequences and 91 bp RNA reads generated) have sensitivity limitations for detecting low, yet specific gene expression in less prevalent cell populations. Overall, genes with lower expression (mainly with a detection at the latest quarter threshold cycle using real time qPCR) were minimally detected at single cell resolution using the current 10X Genomics expression profiling pipeline. However, our approach of combining FACS sorting with cell isolation from the Sost-Cre/Ai9 mice to develop an enriched population of cortical bone cells identified potentially new sets of osteocyte genes which were validated in an independent mesenchymal cell line *in vitro* and in RNA from cortical bone *ex vivo*.

In summary, by combining genomic, transcriptomic, and predictive metabolic profiling approaches, we identified osteolineage genes associated with distinct cell populations of osteoblast precursors, mature osteoblasts and osteocytes. Genes within these three cortical bone cell populations were misregulated in a mouse model of CKD prior to the development of cortical porosity. Thus, our findings support that the molecular events occurring during the bone disease associated with CKD appear early and manifest more widely across the osteolineage cell population.

## Acknowledgements

The authors thank the members of the Indiana University Melvin and Bren Simon Cancer Center Flow Cytometry Resource Facility for their outstanding technical support. This work was supported in part by NIH. The Indiana University Melvin and Bren Simon Comprehensive Cancer Center Flow Cytometry Resource Facility is funded in part by NIH, National Cancer Institute (NCI) grant P30 CA082709 and National Institute of Diabetes and Digestive and Kidney Diseases (NIDDK) grant U54 DK106846. The FCRF is supported in part by NIH instrumentation grant 1S10D012270. The authors would like to acknowledge NIH grants K99-DK129705 (RA), R21-AR059278 (KEW), R01-DK112958 (KEW), P20GM125503 (NIGMS to IN), and the David Weaver Professorship (KEW).

## Data availability

The scRNAseq data have been deposited into the GEO under accession number GSE208152. Raw data for MPC2 cell RNAseq and ATACseq were deposited into the GEO under accession number GSE205792; the data analyzed herein were from undifferentiated (MSCs) and differentiated osteoblasts/osteocytes (OB/OC).

---

**Supplementary Figure 1. A.** Percentage of mitochondrial genes in scRNAseq dataset. **B-F.** Expression density plots indicated cells with high transcription of *Ptprc (Cd45)*, *Cdh5*, *Pdgfra, Mmp13,* and *Spp1*. **G.** The Monocle algorithm divided the cells based upon the original UMAP into three partitions, as indicated by each color. The partition ‘p1’ identified the osteolineage cells, ‘p2’ corresponded to hematopoietic cells and ‘p3’ represented endothelial cells. **H-I.** Examples of trajectory analysis performed on hematopoietic and marrow endothelial cells.

**Supplementary Figure 2. A.** Predictive upstream regulators in Tnc/Mmp13 osteoblast (pre-osteoblasts), osteoblast and osteocyte. **B.** Dmp1 expression in different cell types **C.** scFEA UMAP. **D.** Heatmap indicates the distribution of predicted cell-wise flux of glycolytic, TCA, serine metabolism, fatty acid metabolism and glutamate metabolism relative to pre-osteoblast (Tnc/Mmp13) values. The heatmap uses a column Z-score to show significant differences between pre-osteoblasts (Tnc/Mmp13), osteoblasts, and osteocytes. **E-F.** Ridgeline plots indicate the distribution values of metabolic flux in pre-osteoblasts (P-OB), osteoblasts (OB), and osteocytes (OC). Each ridgeline represents the flux between two metabolites (x-axis) for different cells that are plotted on the y-axis. **G.** Venn diagram shows the number of differentially expressed genes for Tnc_mmp13_osteoblasts vs osteoblasts, and osteocytes vs osteoblasts. **H.** Ridgeline plots indicate the distribution values of metabolic flux in pre-osteoblasts (Tnc/Mmp13), osteoblasts (OB), osteocytes (OC), hematopoietic cells (Hem I and Hem II), and bone marrow endothelial cells (BMEC).

**Supplementary Figure 3. A-F.** Chromatin accessibilities with corresponding gene expression of Spp1 (p = 0.00933495), Col11a1 (p = 1.9363E-05), and Col11a2 (p = 7.2591E-06) in differentiated (OB/OC) *versus* undifferentiated (MSC) cells.

**Supplementary Figure 4. A-C.** Assessment of chromatin accessibilities at *Mmp13* (p = 0.01025011)*, Serpine2* (p = 3.4093E-09), and *Lifr* (p = 1.2547E-07) genomic regions. These genes were detected as highly enriched in pre-osteoblasts. **D-E.** Assessment of chromatin accessibilities at *Serpinf1* (p = 0.00099491), and *Cthrc1* (p = 0.013) genomic loci. These genes were predicted to be highly enriched in osteoblasts. **D-E.** Assessment of chromatin accessibilities at *Ackr3* (p = 2.503E-06)*, Cd109* (p = 0.00179923)*, Ptgis* (p = 6.8707E-06)*, Spns2* (p = 0.00521604), and *Bmp2* (p = 0.00540629) genomic regions. These genes were detected primarily in osteocytes. Top tracks (gray) corresponded to undifferentiated cells (MSC) and lower tracks (black) corresponded to differentiated cells (OB/OC).

**Supplementary Figure 5. Osteocyte and osteoblast cell numbers in cortical and trabecular bone. A.** The image shows osteocytes in the lacunae from femurs that were stained with hematoxylin and eosin. **B.** Osteocyte numbers from cortical bone were counted in the midshaft of bone and normalized to bone area. For counting of osteocytes in trabecular bone, cells were counted in distal femur excluding endocortical surfaces and primary spongiosa, then normalized to trabecular bone area. Osteoblasts were counted in distal femur in the same trabecular region as where osteocytes were counted and normalized to trabecular bone surface.

