## Supplementary figures and images for "Single Cell Cortical Bone Transcriptomics Defines Novel Osteolineage Gene Sets Altered in Chronic Kidney Disease"

### Supplemental Figure 1

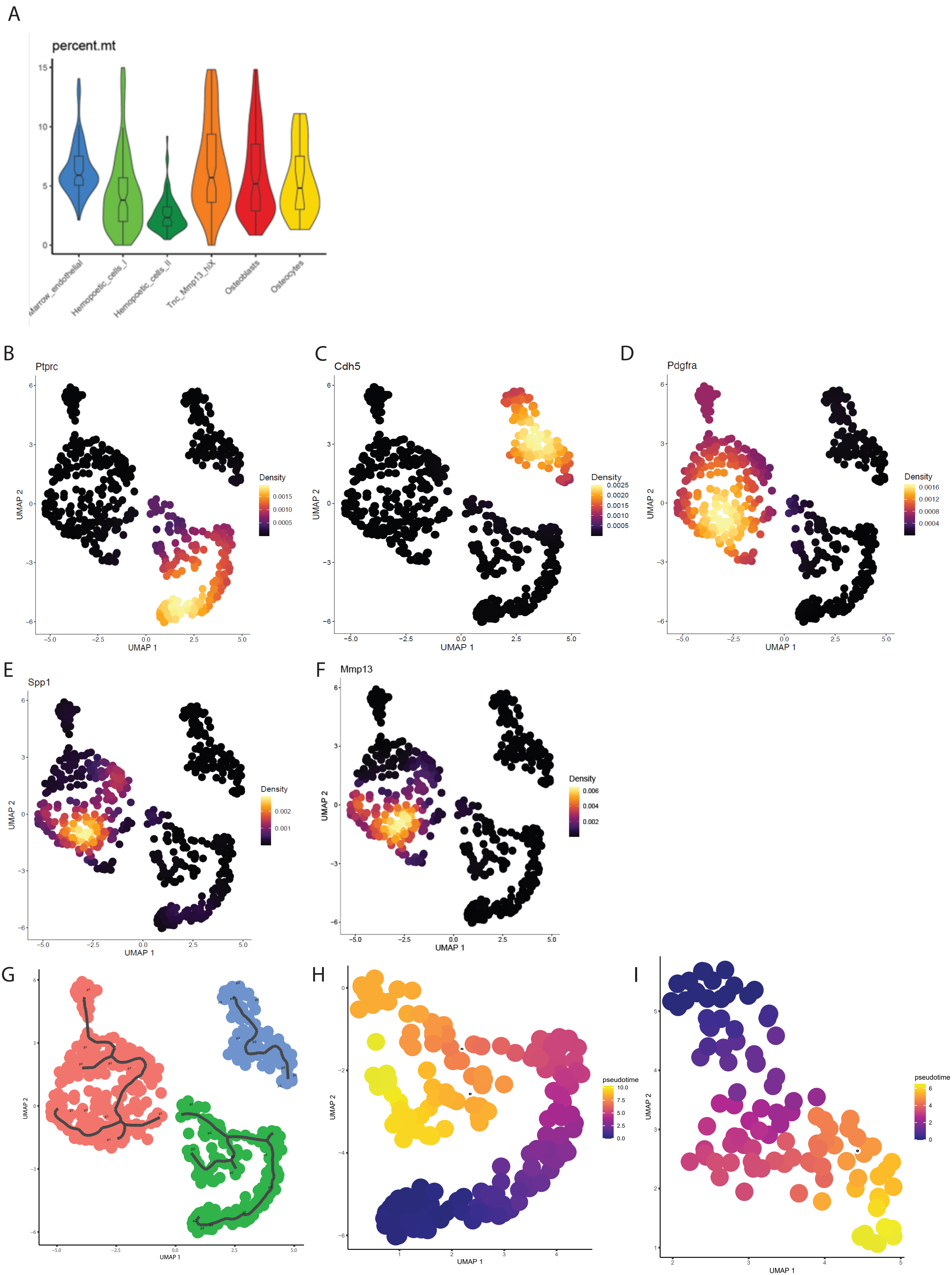

### Supplemental Figure 2

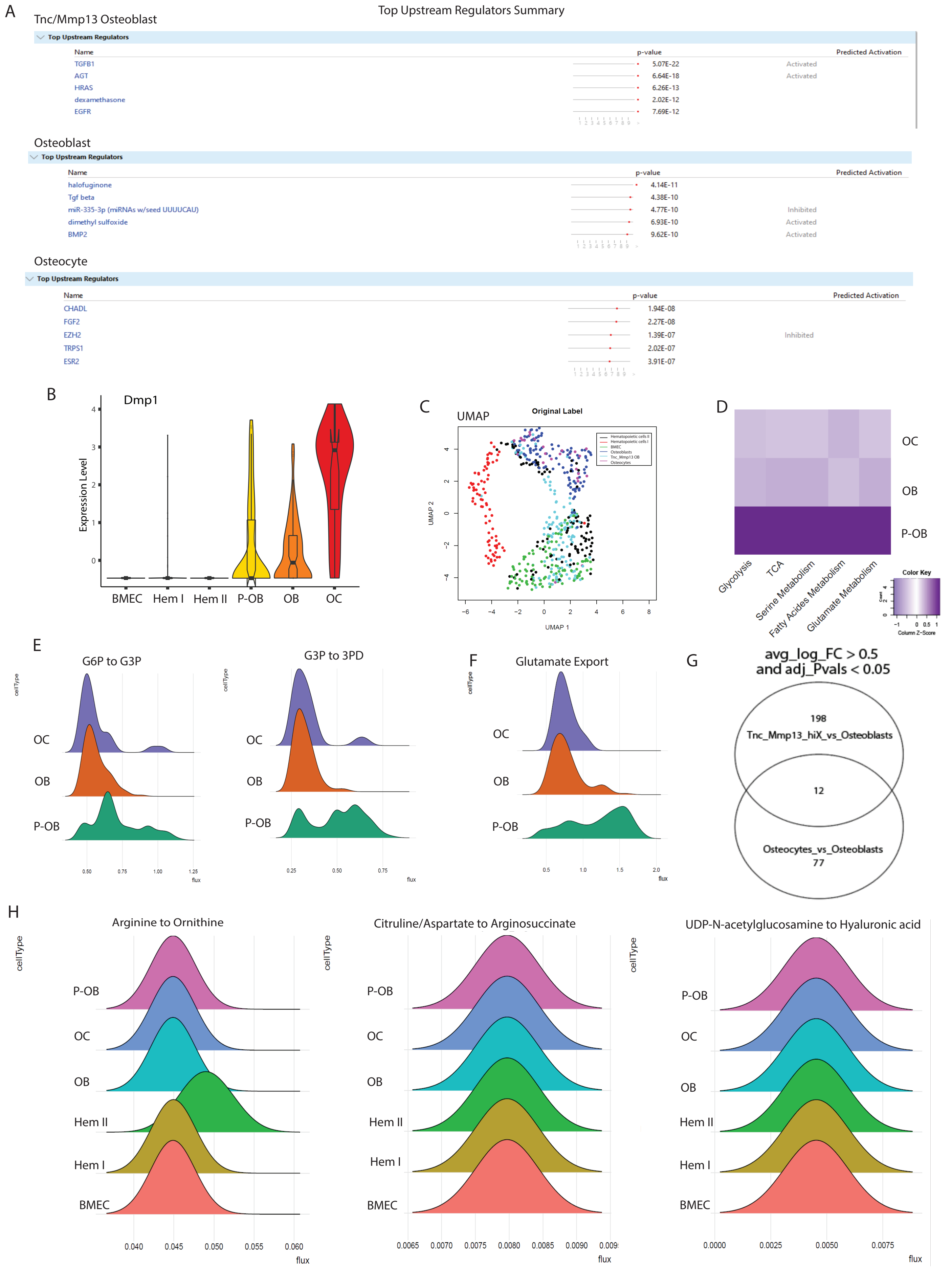

### Supplemental Figure 3

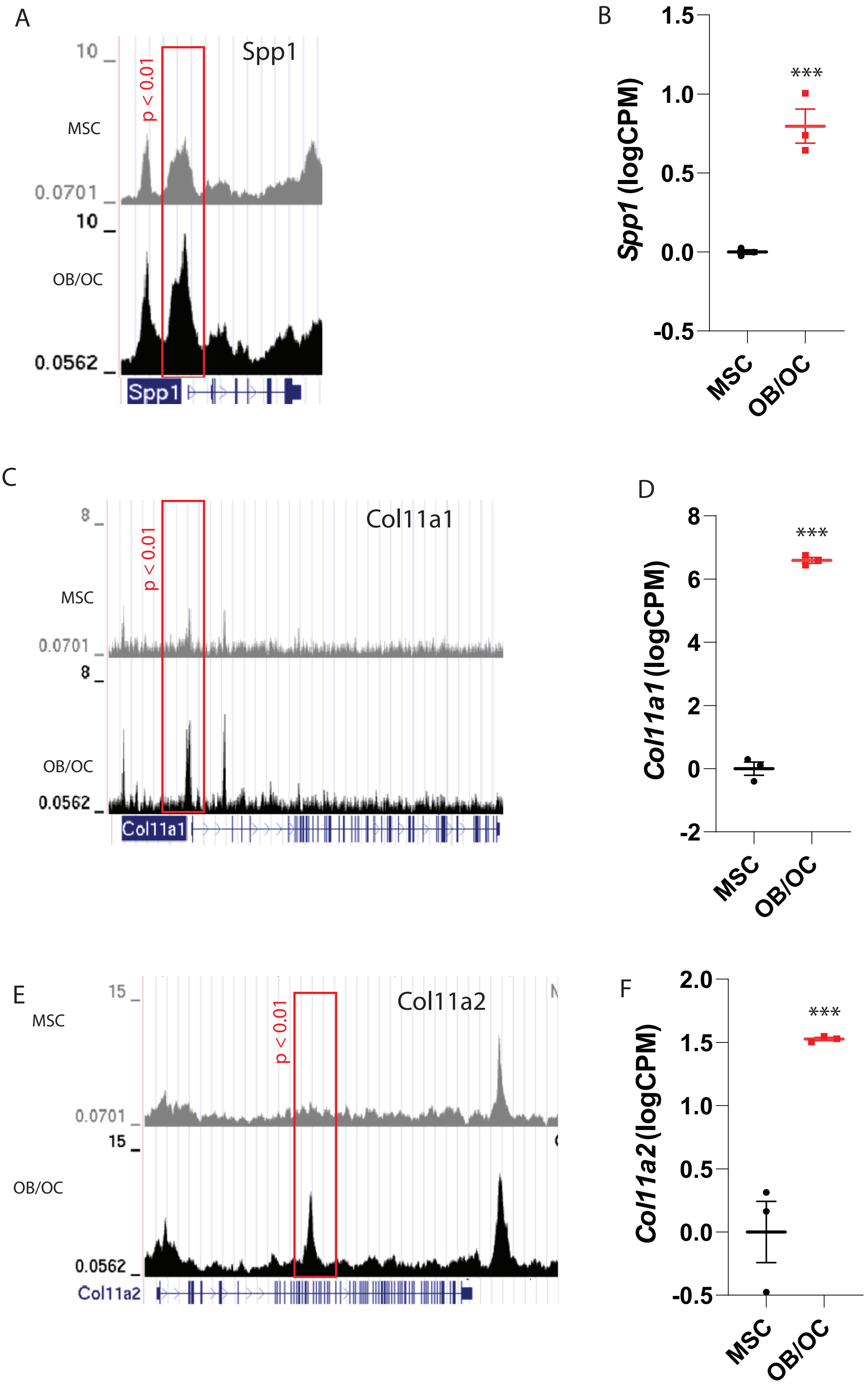

### Supplemental Figure 4

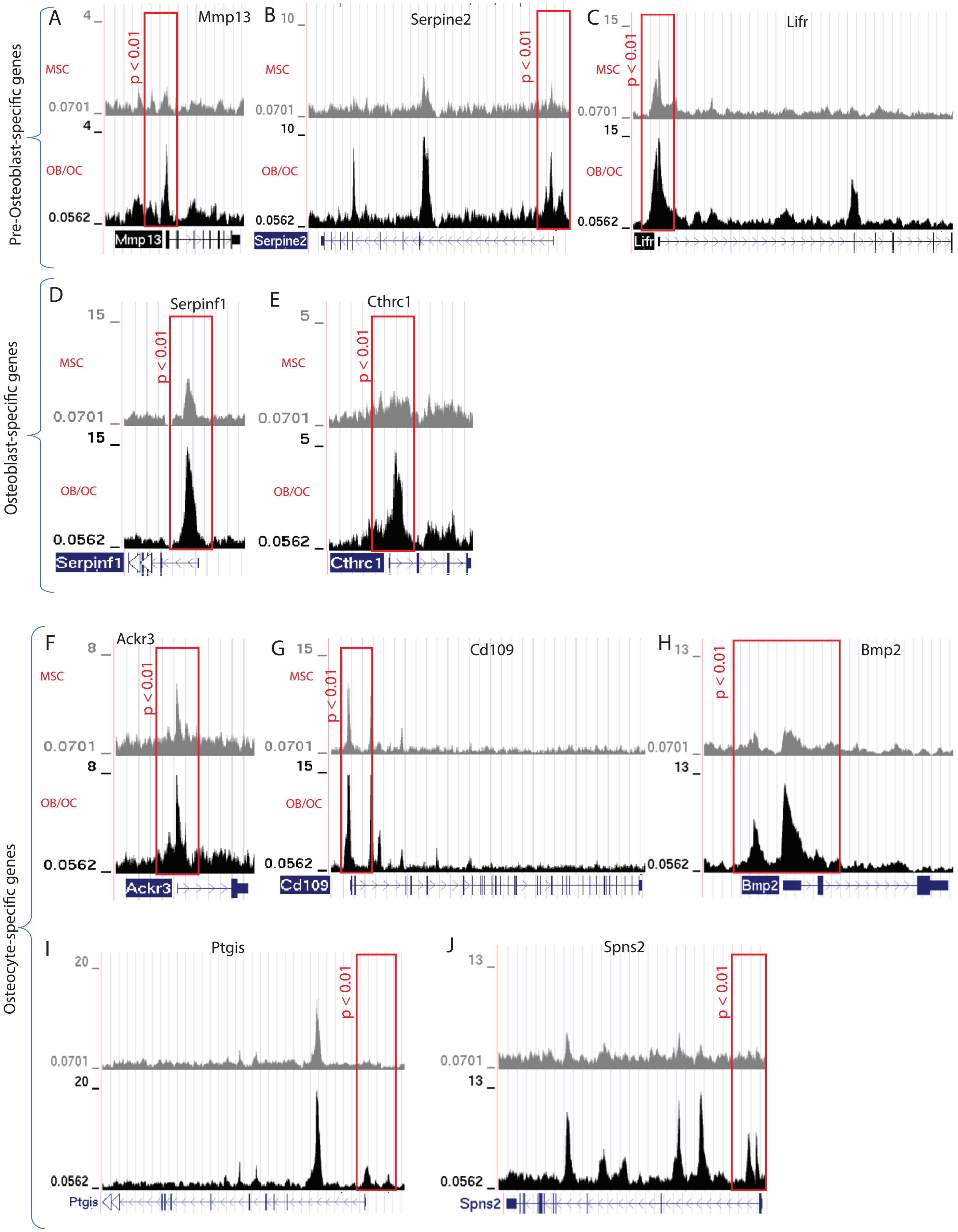

### Supplemental Figure 5

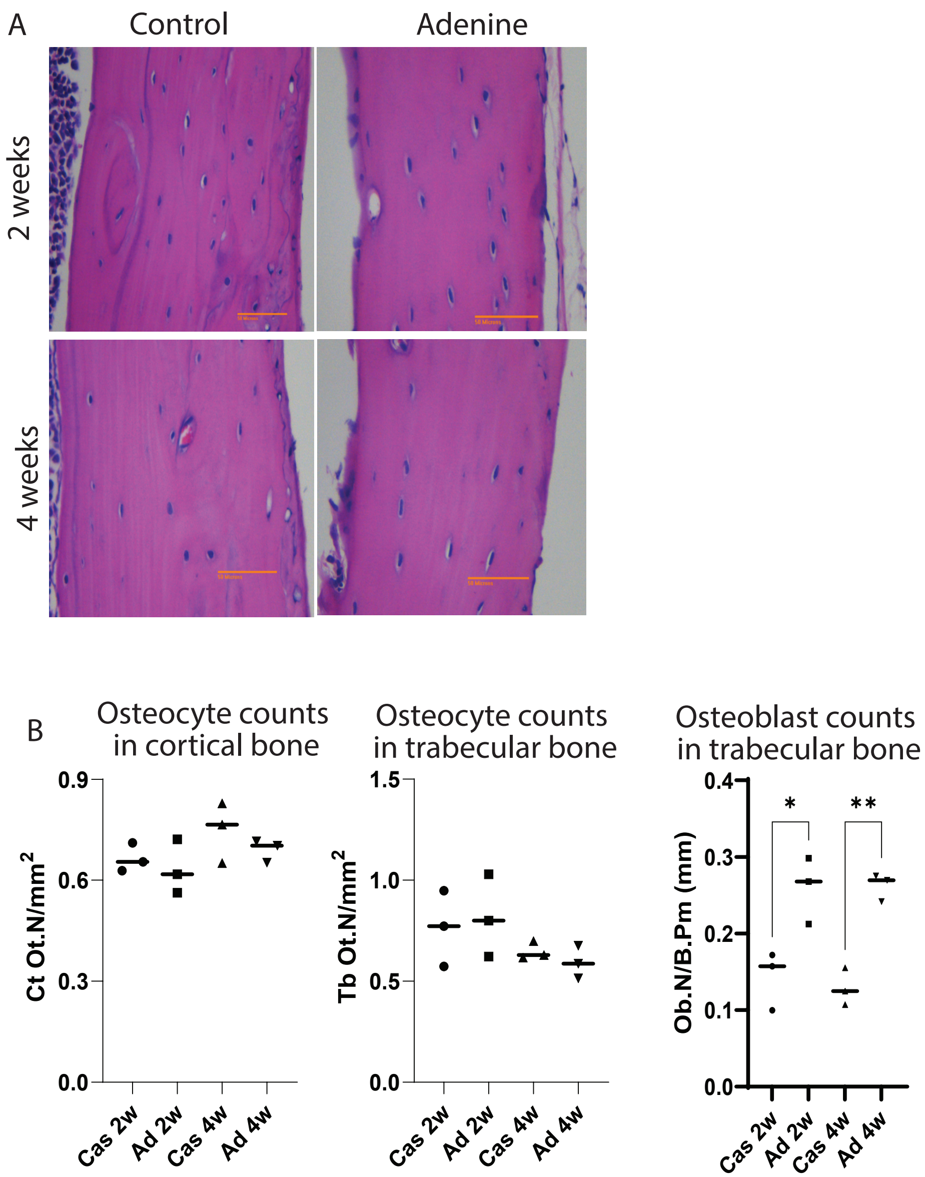
