## Supplementary Table 1 for "Single Cell Cortical Bone Transcriptomics Defines Novel Osteolineage Gene Sets Altered in Chronic Kidney Disease"

**Supplementary Table 1. Osteoblast and osteocyte markers.** Table lists the markers that identified Tnc-Mmp13 osteoblasts, osteoblasts, and osteocytes. The pct.1 is the percentage of cells in the cluster where the gene is detected, whereas pct.2 is the percentage of cells where the gene is detected among all the other cells.

| Tnc_Mmp13 Osteoblasts | | | | | Osteoblasts | | | | | Osteocytes | | | | |
| --- | --- | --- | --- | --- | --- | --- | --- | --- | --- | --- | --- | --- | --- | --- |
| markers | log2FC | pct.1 | pct.2 | p_val_adj | markers | log2FC | pct.1 | pct.2 | p_val_adj | markers | log2FC | pct.1 | pct.2 | p_val_adj |
| Mmp13 | 4.32 | 0.669 | 0.092 | 1.77E-36 | Smpd3 | 3.46 | 0.973 | 0.108 | 1.38E-75 | Phex | 3.91 | 1 | 0.19 | 2.87E-34 |
| Tnc | 3.88 | 0.855 | 0.189 | 6.05E-58 | Bglap | 3.36 | 0.973 | 0.384 | 4.14E-45 | Dmp1 | 3.66 | 0.923 | 0.211 | 2.68E-14 |
| Igfbp5 | 3.59 | 0.589 | 0.109 | 1.09E-24 | Bglap2 | 3.33 | 0.955 | 0.286 | 2.47E-45 | Cd109 | 3.02 | 0.962 | 0.144 | 5.61E-36 |
| Serpine2 | 3.43 | 0.96 | 0.305 | 3.32E-59 | Col11a2 | 3.26 | 1 | 0.165 | 1.17E-69 | Ackr3 | 2.86 | 0.923 | 0.069 | 6.51E-26 |
| Spp1 | 3.10 | 0.79 | 0.364 | 5.47E-29 | Col1a1 | 3.15 | 1 | 0.547 | 2.79E-63 | Ptprz1 | 2.73 | 1 | 0.194 | 3.43E-19 |
| Serping1 | 2.98 | 0.548 | 0.102 | 4.46E-27 | Col1a2 | 2.82 | 1 | 0.499 | 3.35E-55 | Car12 | 2.62 | 0.731 | 0.035 | 4.23E-18 |
| Lifr | 2.84 | 0.798 | 0.421 | 9.60E-43 | Col11a1 | 2.76 | 0.991 | 0.243 | 1.14E-56 | Ramp1 | 2.59 | 0.962 | 0.305 | 2.05E-19 |
| Postn | 2.70 | 0.637 | 0.109 | 8.47E-35 | Serpinf1 | 2.68 | 0.973 | 0.307 | 6.81E-52 | Bambi | 2.55 | 0.923 | 0.386 | 1.48E-14 |
| Cdh11 | 2.57 | 0.766 | 0.243 | 3.61E-52 | Cthrc1 | 2.42 | 0.855 | 0.11 | 7.77E-53 | Dkk1 | 2.54 | 0.923 | 0.067 | 1.17E-19 |
| Gdpd2 | 2.57 | 0.435 | 0.009 | 3.41E-31 | Col5a2 | 2.39 | 1 | 0.378 | 1.24E-55 | Pdgfa | 2.50 | 1 | 0.276 | 5.96E-22 |
| Pdgfrb | 2.53 | 0.589 | 0.054 | 3.39E-42 | Sparc | 2.35 | 1 | 0.65 | 1.05E-51 | Pdpn | 2.29 | 0.885 | 0.067 | 7.27E-20 |
| Gas6 | 2.52 | 0.556 | 0.177 | 6.36E-19 | Ibsp | 2.35 | 0.982 | 0.316 | 3.22E-37 | Wasl | 2.18 | 0.962 | 0.273 | 5.59E-15 |
| Olfml2b | 2.51 | 0.669 | 0.137 | 2.73E-37 | Ifitm5 | 2.29 | 0.873 | 0.126 | 5.79E-48 | Col24a1 | 2.09 | 0.923 | 0.125 | 1.51E-17 |
| Col6a1 | 2.41 | 0.677 | 0.156 | 7.52E-28 | Col5a1 | 2.13 | 0.991 | 0.281 | 2.30E-53 | Ptgis | 2.06 | 1 | 0.307 | 5.67E-13 |
| Col12a1 | 2.37 | 0.637 | 0.118 | 2.49E-29 | Cpz | 2.09 | 0.855 | 0.078 | 3.88E-54 | Bmp2 | 2.05 | 0.846 | 0.113 | 8.36E-15 |
| Lum1 | 2.36 | 0.71 | 0.293 | 2.17E-20 | Ccn1 | 2.08 | 0.718 | 0.256 | 6.98E-16 | Ccn4 | 2.04 | 1 | 0.307 | 6.14E-13 |
| Islr | 2.33 | 0.565 | 0.106 | 6.98E-34 | Cgref1 | 1.96 | 0.827 | 0.101 | 1.93E-49 | Gm41724 | 1.94 | 0.692 | 0.052 | 5.37E-15 |
| Col8a1 | 2.30 | 0.387 | 0.033 | 5.23E-20 | Nupr1 | 1.94 | 0.945 | 0.334 | 1.42E-33 | Spns2 | 1.84 | 0.923 | 0.273 | 7.79E-14 |
| Cfh1 | 2.29 | 0.895 | 0.364 | 3.51E-34 | Fkbp11 | 1.93 | 0.873 | 0.128 | 2.22E-50 | Bmp4 | 1.81 | 0.885 | 0.077 | 1.20E-16 |
| Col6a2 | 2.24 | 0.629 | 0.139 | 1.25E-23 | Rrbp1 | 1.92 | 0.991 | 0.627 | 1.01E-52 | Plpp1 | 1.81 | 1 | 0.246 | 1.07E-11 |
| Loxl1 | 2.20 | 0.5 | 0.043 | 3.96E-32 | Serpinh1 | 1.85 | 1 | 0.558 | 5.32E-43 | Irx5 | 1.78 | 0.962 | 0.069 | 1.22E-21 |
| Wif11 | 2.19 | 0.532 | 0.227 | 1.40E-12 | Timp1 | 1.85 | 0.791 | 0.238 | 1.07E-22 | Myo1b | 1.71 | 0.962 | 0.338 | 7.29E-13 |
| Limch1 | 2.19 | 0.468 | 0.116 | 4.32E-19 | Creb3l1 | 1.84 | 0.891 | 0.185 | 1.24E-44 | Adamts14 | 1.70 | 0.846 | 0.058 | 7.65E-20 |
| Rn18s | 2.18 | 1 | 0.998 | 2.82E-30 | Col22a1 | 1.84 | 0.973 | 0.176 | 2.29E-53 | Cspg4 | 1.58 | 0.923 | 0.171 | 2.58E-11 |
| Cp | 2.18 | 0.661 | 0.132 | 2.62E-33 | Car3 | 1.80 | 0.909 | 0.19 | 1.28E-40 | Ccdc194 | 1.57 | 0.885 | 0.104 | 8.03E-15 |
