## Supplementary Table 2 for "Single Cell Cortical Bone Transcriptomics Defines Novel Osteolineage Gene Sets Altered in Chronic Kidney Disease"

**Supplementary Table 2:** **Jaccard index analysis.** The Jaccard scores were calculated for the four groups of overlapping differentials expressed genes (DEGs) and differential accessible regions (DARs) based on their up- or down-regulated directions: both up, both down, DEG up & DAR down, and DEG down & DAR up.

|  | Cutoffs: FDR < 0.05, \|logFC\| >1 | |
| --- | --- | --- |
|  | Overlap p value | jaccard |
| DAR_Up and DE_Up | 3.25e-35 | 0.0714 |
| DAR_Down and DE_Down | 0.00111 | 0.0296 |
| DAR_Up and DE_Down | 1 | 0.00309 |
| DAR_Down and DE_Up | 1 | 0.00605 |
